# Phosphorylation of α-synuclein fibrils at S129 changes DNAJB1 binding as probed by solid-state NMR

**DOI:** 10.1101/2024.12.19.629500

**Authors:** Sayuri Pacheco, Dhanya S. Reselammal, Qingya Zhang, Ansgar B. Siemer

**Affiliations:** Department of Physiology and Neuroscience, Zilkha Neurogenetic Institute, Keck School of Medicine, University of Southern California, Los Angeles, California, USA

**Keywords:** solid-state NMR, amyloid fibrils, chaperone, post-translational modifications, phosphorylation, J-proteins, α-synuclein, DNAJB1

## Abstract

Amyloid fibrils composed of the protein α-synuclein (aSyn) are implicated in the pathogenesis of synucleinopathies. These pathological fibrils, characterized by their rigid amyloid cores, also feature flexible intrinsically disordered regions (IDRs) that interact with various cellular components. Due to their solvent exposure and flexibility, the N- and C-termini, IDRs of aSyn fibrils, have been used as targets for immunotherapies and serve as binding sites for many chaperones. Chaperones are known to play a vital role in preventing and reversing amyloid formation in neurodegenerative diseases, but how they recognize misfolded proteins is often poorly understood. DNAJB1 is a co-chaperone that is part of a larger complex formed with Hsp70 and Apg2, which collectively disaggregate amyloid fibrils such as those formed by aSyn, tau, or huntingtin. Although the entire chaperone and co-chaperone complex are required for aggregate disassembly, DNAJB1 directly recognizes these amyloid fibrils. More specifically, DNAJB1 preferentially binds to the C-terminus of wild-type aSyn. However, over 90% of aSyn fibrils in Lewy bodies are phosphorylated at S129. Here, we determine the effect of S129 phosphorylation on DNAJB1-aSyn fibril binding using biochemical assays and solid-state NMR. We show that DNAJB1 preferentially binds aSyn pS129 fibrils and that the aSyn binding site of pS129 and wild-type fibrils is different. These results suggest that pS129 might influence chaperone-mediated degradation efficiency.

## Introduction

Synucleinopathies including Parkinson’s disease (PD), dementia with Lewy bodies, and multiple system atrophy are neurodegenerative disorders, characterized by the presence of Lewy bodies, which are primarily composed of α-synuclein (aSyn) fibrils.^1,2^ The aSyn sequence is usually divided into three domains: the N-terminus (residues 1–60), which is of an intermediate flexibility in fibrils;^3,4^ the NAC (non-amyloid-beta component) domain (residues 61–94), which forms the majority of the fibril core;^5^ and the C-terminus (residues 95–140), which is negatively charged and dynamic.^6^ aSyn forms multiple fibril polymorphs in vivo and in-vitro with distinct cytotoxicity and seeding profiles.^7–11^ While the majority of the structural studies have focused on the fibril core, the less-ordered and highly dynamic intrinsically disordered regions (IDRs) framing this core remain understudied. These IDRs form the surface of the fibrils which are exposed to the cellular components, and interact with a range of proteins including chaperones such as protein isomerase 1,^12^ Hsp70,^13^ DNAJB1,^14,15^ and lymphocyte-activation gene 3^16,17^ highlighting the significance of IDRs in neurodegenerative diseases.

Molecular chaperones play a key role in preventing protein misfolding, facilitating protein disaggregation and amyloid disassembly.^18^ Heat shock proteins (Hsp) are a family of highly conserved molecular chaperones,^19^ which were reported to disassemble amyloid fibrils, formed by huntingtin,^20,21^ tau,^22,23^ and aSyn.^24^ Hsp70 has been identified in Lewy bodies,^25^ and upregulation of Hsp70 has been shown to reduce aSyn overexpression in PD mouse models.^26^ Recent studies suggest that the Hsp70 chaperone, along with its co-chaperone DNAJB1 and the nucleotide-exchange factor Apg2, can bind to both aSyn monomers and fibrils, facilitating fibril disaggregation in vitro.^27^ Notably, DNAJB1 specifically recognizes aSyn fibrils and directs Hsp70 to the fibrils to enable disaggregation. DNAJB1 belongs to the family of J-domain proteins that are defined by a conserved J-domain, which interacts with Hsp70. In addition, DNAJB1 has an N-terminal dimerization domain and two C-terminal domains, CTD I and CTD II, which are connected by a linker region and responsible for substrate recognition.^28,29^ Specifically, DNAJB1 binds aSyn as a dimer through its CTD II domain.^30^ Using solution NMR, the specific binding site on aSyn monomers was mapped to residues 123-129 in its C-terminal region,^27^ which is rich in acidic and aromatic residues.

Although various post-translational modifications (PTMs) of aSyn have been identified such as phosphorylation,^31^ acetylation,^32,33^ ubiquitination,^34^ SUMOylation,^35^ nitration,^36^ and O-GlcNAcylation,^37^ studies indicate that about 90% of aggregated aSyn in Lewy bodies is phosphorylated at residue S129.^38 31^ Notably, to form fibrils in vitro phosphorylation of aSyn at S129 (aSyn pS129) must occur post-fibril formation. Monomeric pS129 does not readily aggregate, highlighting a potential regulatory role of this modification in fibril assembly.^15^

Because pS129 is the dominant PTM in Lewy bodies, we wondered what this modification meant for DNAJB1 binding. In addition, we wanted to get high-resolution structural data of DNAJB1 bound aSyn fibrils. In this study, we are using solid-state NMR to directly probe the interaction between DNAJB1 and WT and pS129 aSyn fibrils.

## Methods

### Protein expression, purification, and fibril formation

#### Expression of aSyn (unlabelled)

Wild type aSyn in PRK172 plasmid was transformed into BL21DE3 cells. A single colony was isolated from an ampicillin (AMP) resistant LB-agar plate and inoculated into a 50 mL primary LB culture overnight at 200 rpm 30℃. The next day, 7 mL of primary culture were transferred to 1L of LB medium until OD reached 0.6-0.7. Protein expression was induced with 0.5 M IPTG for 18 hours at 25℃ and 200 rpm. Once expression was completed, cells were harvested at 4000 rpm using the F9-6x1000 LEX rotor for 20 minutes at 21℃. Cell pellets were flashed frozen in liquid nitrogen and stored at -80℃ for further use.

#### Expression of ^15^N, ^13^C aSyn

Uniformly labeled ^13^C, ^15^N WT aSyn was expressed using the approach reported by Marley et al.^39^ Cells were transformed and inoculated the same way as the unlabeled protein. Once OD reached 0.6-0.7, cells were centrifuged at 4,000 rpm using a F9-6x1000 LEX rotor for 20 minutes at 21℃, the supernatant was discarded and the cell pellets were washed in M9 wash buffer. Cells were centrifuged once again at the aforementioned conditions. The cell pellets were resuspended in M9 minimal media containing 100 mg/mL AMP and incubated for 1 hour at 25℃ and 200 rpm. Protein expression was induced with 0.5 M IPTG for 18 hours at 25℃ and 200 rpm. Once expression was completed, cells were harvested at 4000 rpm for 20 minutes at 21℃. Cell pellets were flashed frozen in liquid nitrogen and stored at -80℃ for further use.

#### Purification of aSyn

Protein pellets were flashed frozen with liquid nitrogen and thawed 3 times. Thawed cells were resuspended in 10 mL of lysis buffer (500 mM NaCl, 100 mM Tris, 1 mM EDTA and 10 mM β-mercaptoethanol, pH 8.0). One tablet of Pierce™ Protease Inhibitor, EDTA-free, was dissolved in the protein suspesion followed by incubation on ice for 30 minutes. The mixture was centrifuged at 30,000g for 20 minutes at 4°C. The supernatant containing aSyn was acidified to pH 3.5 with 1 and 6 M HCl and then incubated on ice for 30 minutes. The suspension was then centrifuged at 30,000 g for 20 minutes at 4°C. The supernatant was dialyzed against 4 L of 1% acetic acid in a 6-8 kDa dialysis membrane overnight. The mostly pure protein sample was then filtered through a 0.22 µm PES filter and loaded into a Jupiter 15 µm C4 300 Å, LC column 250 x 4.6 mm for hydrophobic interaction purification using an FPLC. SDS-PAGE was used to confirm the presence and purity of the protein. The pure fractions were pooled and dialyzed against 50 mM phosphate buffer pH 7.4 with 0.1 mM EDTA and 0.02% NaN_3_ for further use.

### Fibril formation

After dialyzing in 50 mM phosphate buffer pH 7.4, 0.1 mM EDTA, and 0.02% NaN_3_, monomeric aSyn was diluted to 100 µM. For fibrillization, aliquots of 500 µl at 100 µM protein concentration were placed in 1.5 mL low binding tubes with one 1 mm glass bead. Tubes were then sealed with parafilm to prevent evaporation and placed in MATRIX Orbital Delta F1.5 thermomixer, shaking at 300 rpm at 37℃ for 2 weeks.

### Phosphorylation of aSyn fibrils and Western blot analysis

aSyn fibrils were ultra centrifuged twice at 65,000 rpm, 21℃, for 1 hour using a Beckman TLA 100.3 to remove the fibrillization buffer and then fibrils were washed with nuclease free water. The fibrils were resuspended in phosphorylation buffer (20 mM HEPES, 10 mM MgCl_2_, 1.09 mM ATP, 2 mM DTT, pH 7.4) and briefly bath sonicated. PLK2 phosphorylation enzyme was then added to the fibrils (1 µg of enzyme per 1.44 mg of protein). The reaction was carried out by incubating the samples at 37℃ with mild shaking for 36 hours. The reaction was stopped by brief bath sonication and freezing samples at -80℃. Phosphorylation of the fibrils was confirmed using western blot with the Ep1536Y antibody (ab51253) against phosphorylated S129 aSyn.

### Expression and purification of DNAJB1

Plasmid containing DNAJB1 construct having N-terminal His-SUMO tag was transformed into chemically competent BL21(DE3) cells. Starter cultures containing 50 mg/mL kanamycin were grown for 14–16 hours at 30°C. The culture was then expanded into LB medium and grown at 37°C to an OD_600_ of 0.6-0.8, after which protein expression was induced using IPTG at a final concentration of 1 mM. The cell culture was incubated at 20°C, 140 rpm for 20 hours. Cells were harvested by centrifugation at 6000 g for 20 minutes and stored at -80°C for further use. For purification, cells were resuspended in lysis buffer composed of 30 mM Hepes pH 7.4, 500 mM KCl, 5 mM MgCl_2_, 30 mM imidazole, and 10% glycerol containing phenylmethylsulfonyl fluoride (PMSF), Pierce Protease Inhibitor EDTA-free, DNase I, and β-mercaptoethanol. Cells were then sonicated using a QSonic Ultrasonic Sonicator and cell lysate was centrifuged for 30 minutes at 20,000 rpm using an F21-8x50y rotor. The supernatant was added to His60 Ni Superflow resin and incubated on a shaker at 4°C for 1 hour. The column was then washed with high salt buffer (30 mM Hepes pH 7.4, 1 M potassium acetate (KAc), 5 mM MgCl_2_, 25 mM imidazole, and 10% glycerol) followed by low salt buffer (30 mM Hepes pH 7.4, 50 mM KAc, 5 mM MgCl_2_, 25 mM imidazole, and 10% glycerol) both containing β-mercaptoethanol. The protein was then eluted with a buffer containing 30 mM Hepes pH 7.4, 100 mM KAc, 5 mM MgCl_2_, and 300 mM imidazole and β-mercaptoethanol. The eluted protein was combined with Ulp1 protease at a ratio of 400 μg of protease per 1 mL of substrate and dialyzed against 30 mM Hepes pH 7.4, 100 mM KAc, 5 mM MgCl_2_, and 10% glycerol with β-mercaptoethanol. After dialysis the sample was centrifuged for 20 minutes at 4,000 rpm using an Eppendorf A-4-44 rotor and the supernatant was then added to equilibrated His60 Ni Superflow column. The sample in the column was incubated at 4°C with mild shaking for 20 minutes, after which the flowthrough, containing the cleaved chaperone, was collected.

### Sedimentation assay

The phosphorylated fibrils were ultra centrifuged at 179,343 rcf to remove the PLK2 enzyme and the phosphorylation buffer components. For the DNAJB1 binding study, the aSyn WT, and aSyn pS129 fibrils were incubated with DNAJB1 at a ratio of 5:1 (aSyn:DNAJB1) in 30 mM HEPES buffer containing 0.02% NaN_3_ at pH 7.4, overnight at room temperature with mild shaking. The samples were ultracentrifuged at 179,343 rcf to pellet down fibrils and bound DNAJB1 leaving the unbound DNAJB1 and aSyn monomers in the supernatant. The amount of unbound DNAJB1 in the supernatant was estimated using SDS-PAGE. The morphology of the bound fibrils was analyzed using electron microscopy.

### Electron microscopy of bound and unbound fibrils

Electron micrograph grids were prepared by loading 10 µL of the fibril sample onto Formvar-carbon-coated copper mesh grids (Electron Microscopy Sciences), which were then incubated for 5 minutes. The excess sample was removed from the grids using filter paper. The grids were stained three times with 10 µL of 1% uranyl acetate, with each incubation lasting 5 minutes. After each incubation, excess uranyl acetate was blotted off with filter paper. The grids were then washed with filtered, deionized H_2_O, blotted dry with filter paper, and left to dry overnight.

The electron micrographs were taken on a Jeol JEM-1400 electron microscope, which was operating at 100 kV and equipped with a Gatan Orius CCD camera. Micrographs were captured at various magnifications as indicated.

### Solid-state NMR spectroscopy

All solid-state NMR experiments were conducted on an Agilent DD2 600 MHz solid-state NMR spectrometer equipped with a T3 1.6 mm probe, operating at 25 kHz magic angle spinning (MAS), unless otherwise specified, and a set temperature of 0°C. ^1^H, ^13^C, and ^15^N hard radio frequency (rf) pulses had amplitudes of 200 kHz, 100 kHz, and 50 kHz, respectively. ^1^H–^13^C cross-polarization (CP) experiments utilized the Hartmann-Hahn matching condition at 60 kHz and 85 kHz for ^13^C and ^1^H, respectively, with a 10% amplitude ramp.

One dimensional CP and refocused INEPT (insensitive nuclei enhanced by polarization transfer) experiments were recorded with a spectral width of 50 kHz and 1000 complex time domain points were recorded. A recycle delay of 3 seconds was used to optimize relaxation recovery, with a total of 1024 scans. 140 kHz two-pulse phase modulation (TPPM) decoupling was applied during acquisition.

^13^C-^13^C DREAM^40^ (dipolar recoupling enhanced by amplitude modulation) spectra were recorded at a MAS rate of 30 kHz, with spectral widths of 50 kHz in both dimensions. One thousand complex points were recorded in the direct and 1000 real points (TPPI) were recorded in the indirect dimension. A recycle delay of 3 seconds was applied, and 40 scans were accumulated for each t_1_ increment. The DREAM mixing time was 3.5 ms.

^1^H-^15^N 2D HSQC spectra were acquired with spectral width of 10 kHz and 500 complex points in the direct ^1^H dimension and 3 kHz spectral width and 60 complex points for the indirect ^15^N dimension. 128 scans per t_1_ increment were acquired. The pulse sequence included a BIRD filter with a water relaxation delay of either 320 or 350 ms, which served to suppress residual water signal.^41,42^ WALTZ-16 decoupling with a rf-field strength of 10 kHz was applied on ^15^N.

The HNCA and HNcoCA spectra were recorded as described previously^43^ using non-uniform sampling (NUS) with a sampling density of 60% in both ^13^C and ^15^N dimensions. A total of 32 t_1_ increments were acquired and the NUS data were reconstructed using the IST algorithm.^44^

2D ^1^H-^13^C INEPT-HETCOR spectra of the samples were acquired with 64 scans using a refocused INEPT transfer. The direct ^13^C dimension had a spectral width of 50 kHz with 1000 points; and the indirect ^1^H dimension had a spectral width of 10 kHz with 128 points. Five kHz WALTZ-16 decoupling was applied on ^1^H during acquisition. 2D ^13^C-^13^C INEPT-TOCSY^45^ experiments were conducted with 128 scans, using a FLOPSY8 spin lock sequence for a mixing time of 17 ms. The direct dimension had 50 kHz spectral width with 1000 complex points, and the indirect dimension had 10 kHz spectral width with 75 complex points.

Spectra were processed using nmrPipe version 10.9 and plotted with nmrglue version 0.9 using in-house python scripts. Resonance assignments and peak integration were performed using CARA software.^46^

## Results

### DNAJB1 preferably binds aSyn phosphorylated at S129

Because phosphorylation at S129 is the most common PTM of aSyn fibrils, we investigated whether this modification influences DNAJB1’s recognition of aSyn fibrils. aSyn WT fibrils were phosphorylated at S129 using PLK2, as confirmed by western blot (Figure S1 A). We first quantified the binding affinity of WT and pS129 aSyn fibrils to DNAJB1 using a sedimentation assay. In this assay, WT and pS129 aSyn fibrils were incubated overnight at room temperature with DNAJB1 at a 5:1 molar ratio (aSyn:DNAJB1) (Figure 1A). Afterwards, fibrils were sedimented using centrifugation and the supernatant was analyzed by SDS-PAGE. The SDS-PAGE showed that DNAJB1 was pulled down by both WT and pS129 fibrils, indicated by a decrease in DNAJB1 band intensity. The DNAJB1 band in the supernatant was weaker in the pS129 fibrils compared to the WT sample, suggesting that DNAJB1 is preferentially pulled down by pS129 fibrils (Figure 1B). We used densitometry to quantify the band intensities of unbound DNAJB1 in the supernatant. On average, aSyn pS129 fibrils pulled down 2.7 times more DNAJB1 than aSyn WT fibrils (Figure 1C). We also confirmed the presence of DNAJB1 in the fibril pellet of both fibril types by solubilizing the pellets in 1% DMSO using SDS-PAGE (Figure S1 B)

**Figure 1:**
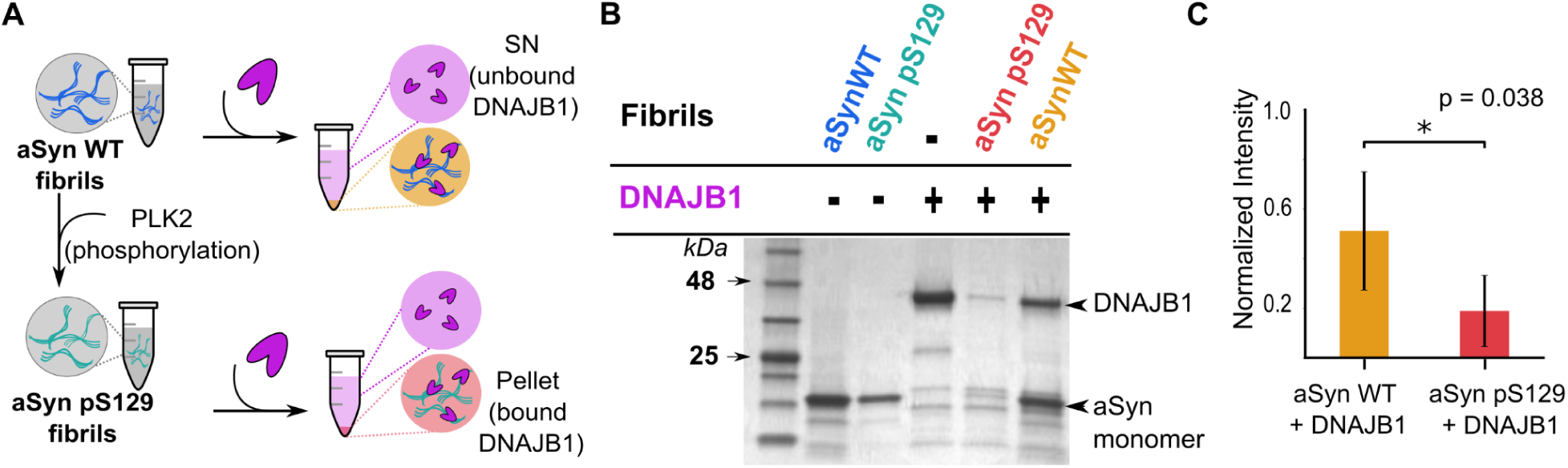
DNAJB1 is preferentially pulled down by aSyn pS129 fibrils compared to WT fibrils. A) Schematic of sedimentation assay for aSyn WT and pS129 fibrils incubated with DNAJB1. WT fibrils were phosphorylated at S129 using PLK2 enzyme. Both fibril types were incubated with DNAJB1 at a ratio of 5:1 (aSyn:DNAJB1). B) Representative SDS-PAGE of supernatants from sedimentation assay. C) Bar graphs showing densitometric quantification of unbound DNAJB1 in the supernatant across five independent experiments.

### DNAJB1 binding leads to fragmentation of aSyn fibrils

The morphology of the aSyn fibrils upon binding to DNAJB1 was analyzed using electron microscopy (EM). Following the sedimentation assay, EM grids were prepared for the four fibril pellets: aSyn WT, aSyn WT with DNAJB1, aSyn pS129 and aSyn pS129 with DNAJB1. Both unbound fibrils (aSyn WT and aSyn pS129) showed previously reported aSyn fibril morphology (Figure 2). The average fibril widths of unbound WT and pS129 fibrils were 11.21 ± 2.04 nm and 9.15 ± 1.94 nm, respectively. The pS129 fibrils had a narrower width distribution as compared to WT fibrils (Figure 2). These fibrils were slightly bundled and had occasional lateral association. While the bound WT and pS129 samples had average fibril widths of 10.59 ± 2.58 nm and 9.25 ± 2.20 nm, respectively. The histograms reveal no significant differences in fibril width upon the addition of DNAJB1, indicating that DNAJB1 does not alter the average fibril width.

**Figure 2:**
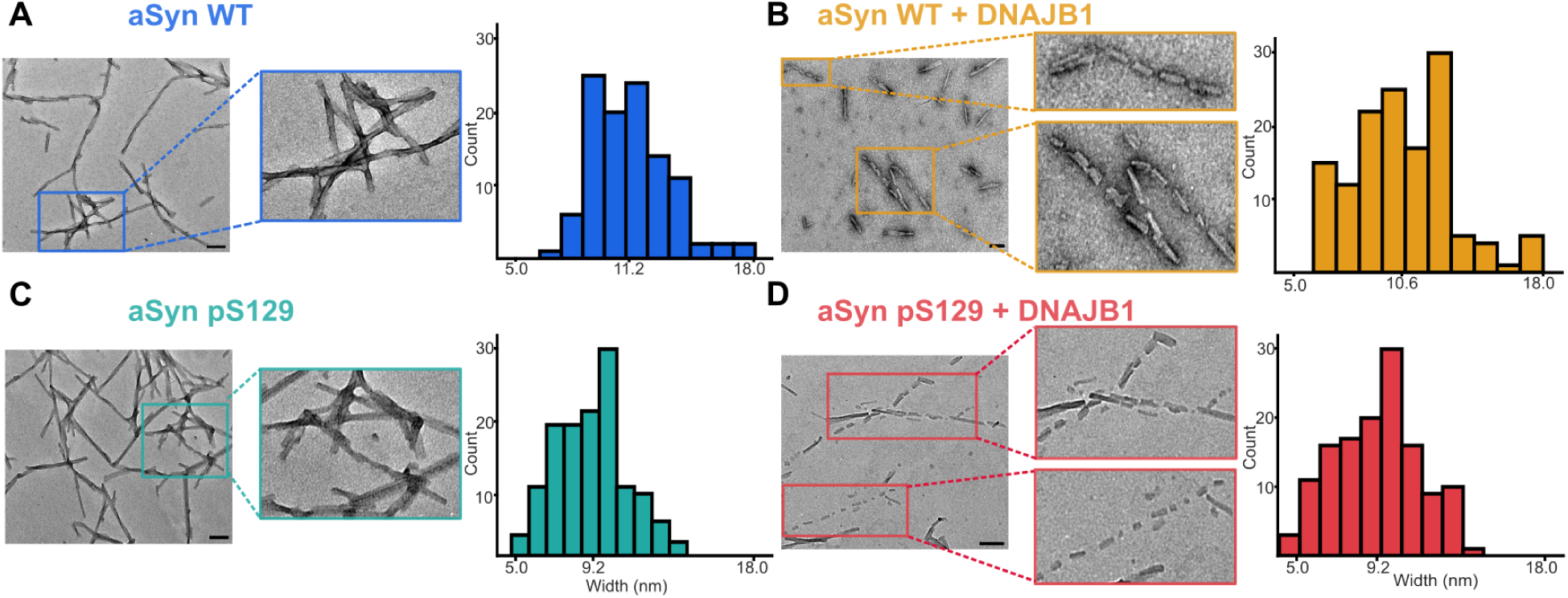
EM images show aSyn fibril fragmentation in the presence of DNAJB1. Scale bars: 100 nm. In the absence of DNAJB1, WT aSyn fibrils (cobalt blue) and phosphorylated aSyn pS129 fibrils (teal) exhibit typical morphologies, with average fibril widths of 11.21 ± 2.04 nm and 9.15 ± 1.94 nm, respectively. Fibril width measurements were obtained manually from EM images. Zoomed-in regions highlight intact fibrils with characteristic lengths and bundled arrangements. Upon incubation with DNAJB1, both fibril types appear fragmented in the micrographs. aSyn WT fibrils in the presence of DNAJB1 have an average width of 10.59 ± 2.58 nm, while pS129 fibrils with DNAJB1 have an average width of 9.25 ± 2.20 nm. The resulting fibril fragments are shorter and clustered, suggesting that the fibrils are more susceptible to mechanical stress. The histograms reveal no significant differences in fibril width upon the addition of DNAJB1, indicating that DNAJB1 does not alter the average fibril width under the conditions tested.

Interestingly, the presence of DNAJB1 induced apparent fibril fragmentation. EM images revealed fragmented fibrils resembling “sausage-like” segments, with breaks occurring at somewhat regular intervals. The fragments were closely spaced but not perfectly aligned. Because we did not observe substantial differences in width distributions of fibrils incubated with DNAJB1, we speculate that the fragmentation arises from mechanical forces during EM grid preparation. These observations suggest that DNAJB1 either directly fragments the fibrils or renders them more susceptible to breakage on the EM grids.

### DNAJB1 binding does not affect the fibril core structure of aSyn

To characterize the DNAJB1-fibril interaction at the molecular level, we prepared uniformly ^13^C,^15^N labeled aSyn fibrils. Four samples were prepared for solid-state NMR experiments, namely aSyn WT, aSyn WT with DNAJB1, aSyn pS129, and pS129 with DNAJB1. Initially, we recorded 1D ^13^C CP and INEPT experiments to assess overall dynamics. The 1D CP overlays of bound and unbound WT and pS129 fibrils showed nearly-perfect overlap, while we observed clear differences in the INEPT spectra (Figure S2). Specifically, the aliphatic region of the INEPT spectra showed evident changes, indicating differences in the dynamic regions when bound to DNAJB1. A comparison of the INEPT/CP ratio between the samples showed a decrease in this ratio with phosphorylation, but an increase in both DNAJB1 bound samples. These results suggest a decrease in dynamics with phosphorylation but an increase in dynamics upon DNAJB1 binding (Figure S3).

To check whether or not DNAJB1 interacts with the fibril core, we recorded 2D ^13^C-^13^C DREAM experiments on all four fibril samples. DREAM spectra detect only rigid regions, such as the fibril core, whose dipolar couplings are not averaged out by molecular motions.^47^ The 2D DREAM spectra of WT and pS129 fibrils showed no significant changes, confirming that phosphorylation at S129 does not affect the fibril core. This result aligns with the fact that phosphorylation occurs after fibril formation and is located in the dynamic C-terminus. Overlaid DREAM spectra of bound and unbound WT and pS129 fibrils (Figure 3) were nearly identical, indicating that DNAJB1 binding does not affect the fibril core. While minor differences in sensitivity were observed, the cross-peaks in the bound and unbound spectra matched well. These data suggest that the interaction between aSyn fibrils and DNAJB1 does not involve the fibril core.

**Figure 3:**
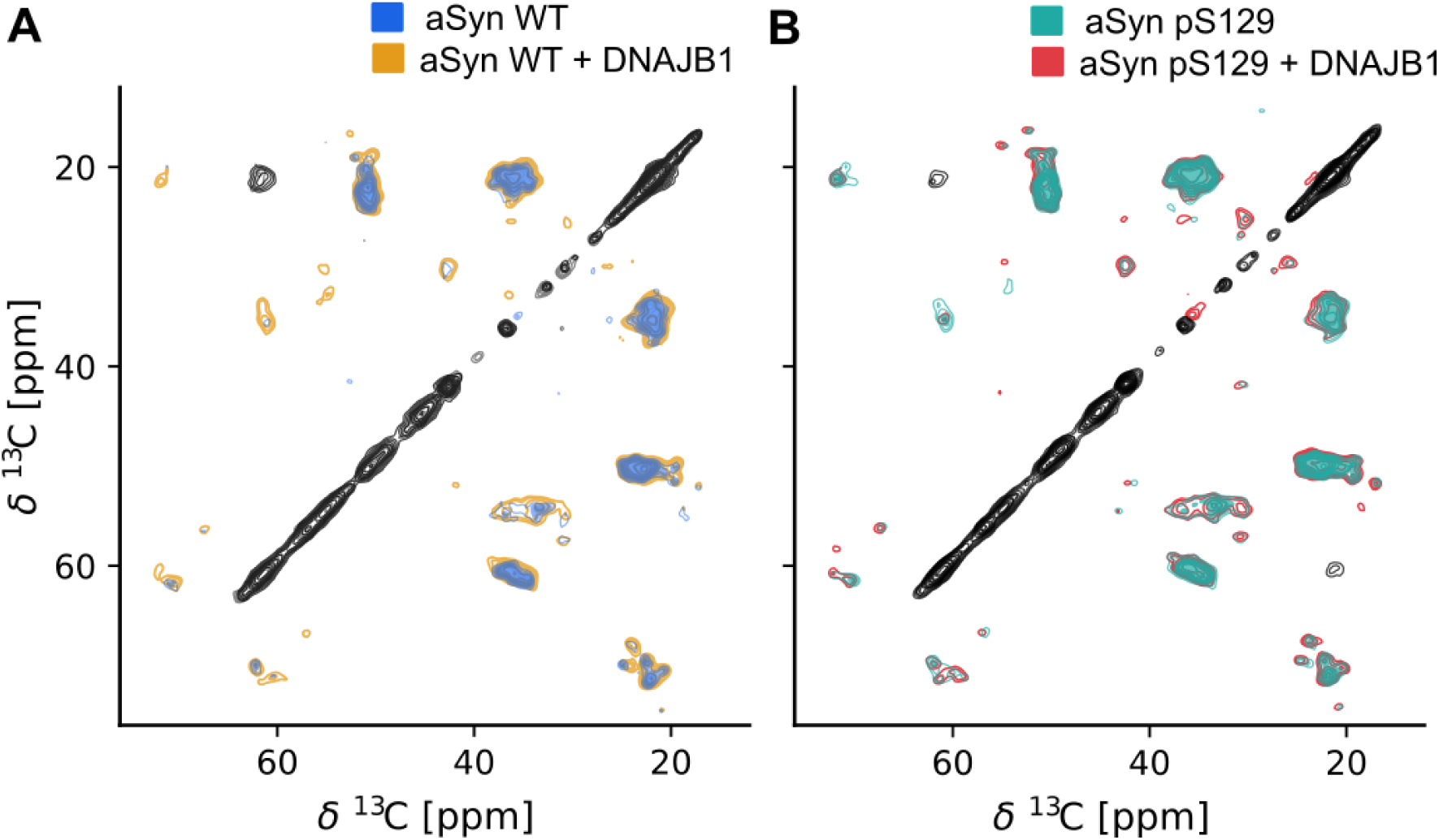
DNAJB1 binding does not affect the fibril core of aSyn fibrils. A) Overlay of ^13^C-^13^C DREAM spectra of aSyn WT fibrils alone and aSyn WT fibrils bound to DNAJB1. B) Same as A but with aSyn pS129 fibrils. The spectra of both the bound fibrils showed a perfect overlap with their spectra of the unbound fibrils suggesting that DNAJB1 binding is not affecting the fibril core of both fibril types.

### DNAJB1 binds to the dynamic C-terminus of both aSyn WT and aSyn pS129 fibrils, with different mode of binding

To characterize the dynamic residues of WT and pS129 fibrils, we recorded 2D ^1^H-^15^N HSQC spectra (Figure 4 A, C). To assign the HSQC of WT aSyn fibrils, we recorded liquid-state NMR type 2D ^13^C- ^1^H INEPT-HETCOR, and 3D HNCA and HNcoCA NMR experiments (Figure S4, S5). Using these data, we assigned twenty six residues from the highly dynamic C-terminus of the fibril. Specifically, we were able to identify most of the non-proline residues starting from A107 until A140, the last residue of the C-terminus. The resolved peaks had an average linewidth of 26 Hz in the proton dimension. The HSQC spectrum of the bound form of the aSyn WT showed clear chemical shift changes (Figure 4A, S6 A). A set of residues in the spectrum showed either a drastic decrease (Q109, G111, E131, G132, Q134, and D135) in intensity or were absent (A107, L113, N122, E123, E126, M127, S129, and E130) from the spectrum in the presence of DNAJB1. The extreme C-terminal peak A140 doubled in the presence of DNAJB1. Additionally, we observed six new peaks in the bound WT HSQC, which are highlighted in purple in Figure 4A. Though we couldn’t unambiguously assign the new peaks, we were able to determine the residue types. We confirmed amino acid type assignments with 2D ^13^C- ^1^H INEPT-HETCOR and ^13^C-^13^C TOCSY spectra (Figure 5A, S7). The five new peaks corresponded to glutamic acids (labeled E_B1_ and E_B2_), two valines (labeled V_B1_ and V_B2_) (Figure 4A, 5A), and tyrosine (labeled Y_B1_in Figure 5A). Furthermore, the HETCOR showed new correlations from a glycine (G_B1_) and alanine (A_B1_) that lacked corresponding HNCA correlation, leaving their assignments ambiguous. The new peaks observed in the HSQC of the WT-bound sample had corresponding ^1^H-^13^C correlations in the HETCOR spectrum (Figure 5A), and showed complete spin system correlations in the TOCSY (Figure S7), enabling us to confirm the residue types. The E_B1_ peak was preceded by a proline residue, as evident from the HNCA spectrum of the bound sample (Figure S8). Notably, aSyn contains only five prolines, all located in the C-terminus (P108, P117, P120, P128, and P138). Additionally, we observed new correlations arising from a proline (linked to the preceding peak in the HNCA) in both the HETCOR and TOCSY spectra (Figure S7). These observations indicate that one of the new peaks, E_B1_, corresponds to the C-terminal residue E139 (P138-E139) in an alternate conformation, as it is preceded by a proline (Figure S8). The new peak E_B2_ could correspond to E137 in an alternate conformation, as it is preceded by a tyrosine (Figure S8). In the bound HETCOR and TOCSY spectra (Figure 5, S7), we observed peak doubling of tyrosine resonances, which contrasts with the unbound sample where a single intense resonance was present for the group of tyrosines. It is likely that the residues associated with the E_B1_ and E_B2_ peaks are E139 and E137, located within the extreme C-terminal region Y136-E137-P138-E139.

**Figure 4:**
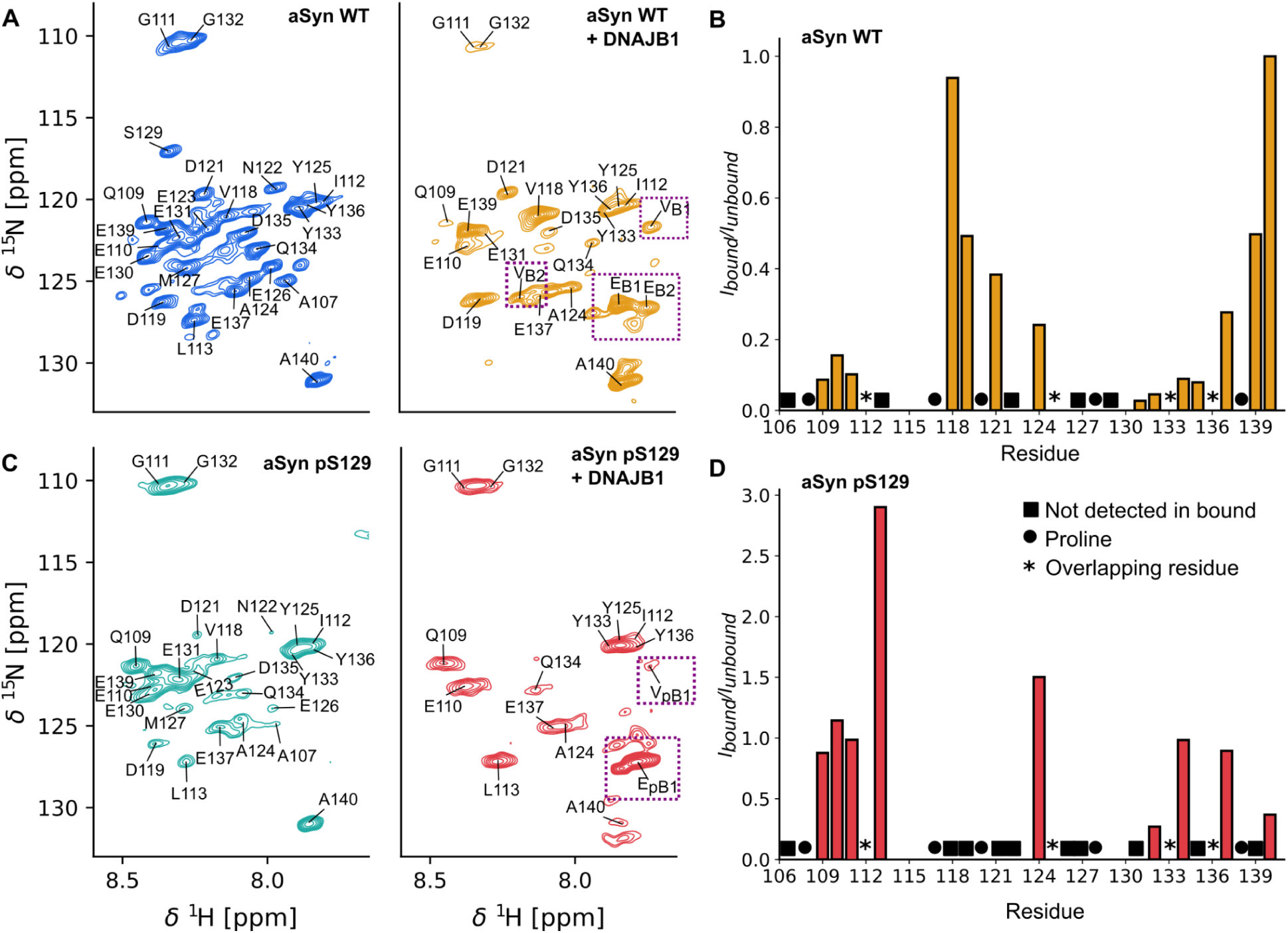
DNAJB1 binding occurs at distinct C-terminal residues in WT and pS129 fibrils. A) HSQC spectrum of WT fibrils (cobalt blue) and WT fibrils with DNAJB1 (gold). The absence of several residues [L113, N122, S129, M127, E130, Q134] in the presence of DNAJB1 indicate clear chemical shift perturbations in the C-terminus of the fibrils. The new peaks that arose in the presence of DNAJB1 are boxed in purple. B). Intensity plot showing effect of DNAJB1 binding with aSyn WT fibrils. I_bound_/I_unbound_ against C-terminal residues (107-140) is shown. C) HSQC spectrum of unbound pS129 fibrils (teal) and pS129 fibrils bound to DNAJB1 (crimson). In comparison to bound WT fibrils, many more peaks are absent in the bound pS129 aSyn fibrils. The most pronounced and isolated peaks remaining in the presence of DNAJB1 are Q109, A107, L113, E130, and G132. Similarly, new peaks arising in the presence of DNAJB1 are boxed around in purple. D). Intensity plot showing effect of DNAJB1 binding with aSyn pS129 fibrils. I_bound_/I_unbound_ vs C-terminal residues (107-140) is shown. Unlike the DNAJB1 binding region in aSyn WT fibrils (Figure 3B), where the most affected residues are located in the middle of the C-terminus (125-138), DNAJB1 binding in aSyn pS129 fibrils affects both the middle and extreme C-terminal regions (Figure 3D).

**Figure 5:**
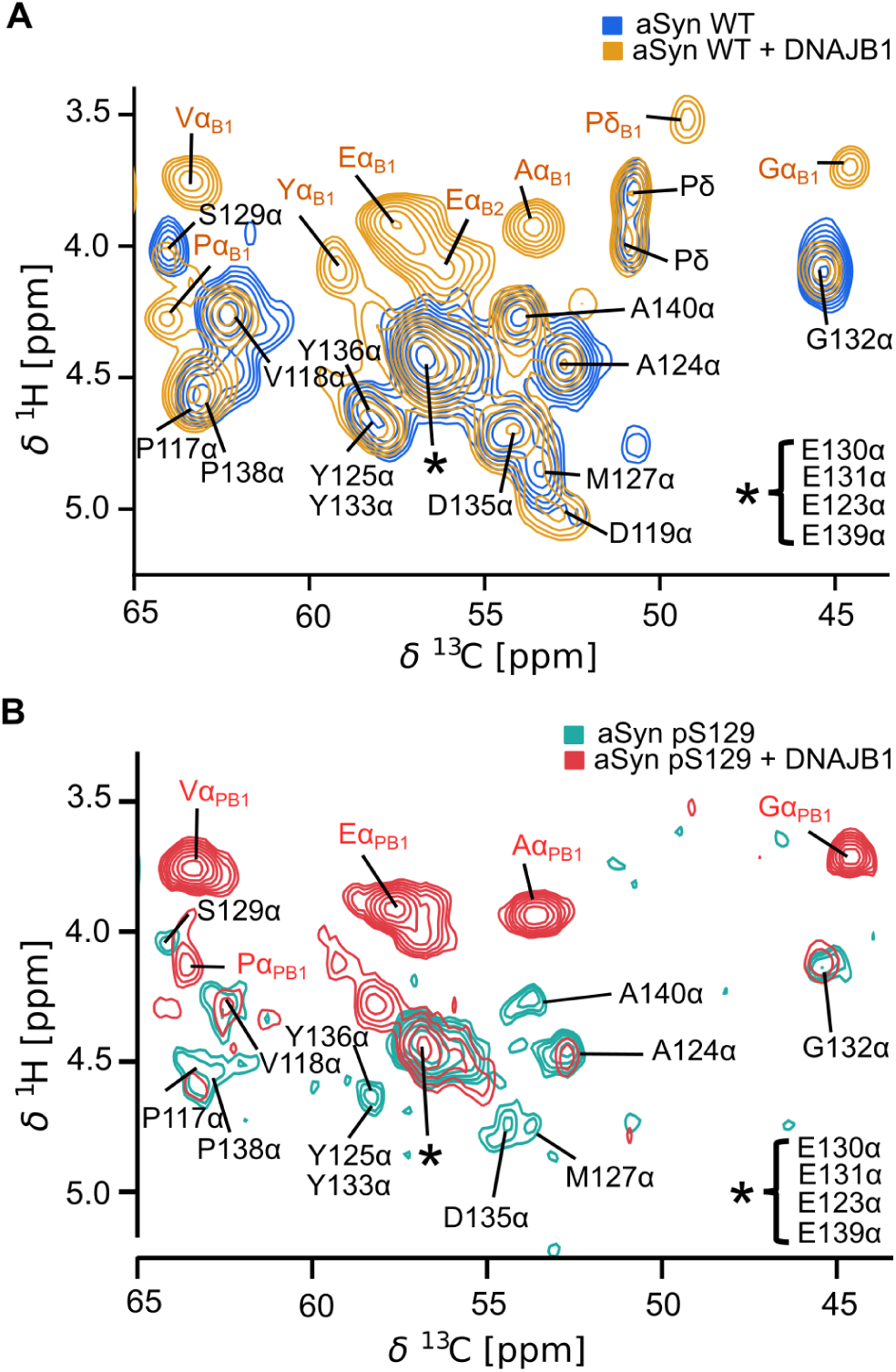
INEPT-HETCOR spectra reveal chemical shift perturbations at the dynamic C-terminus, consistent with the HSQC analysis. A) Overlaid 2D ^1^H-^13^C INEPT-HETCOR spectra of aSyn WT without DNAJB1 (blue) and with DNAJB1 (gold). Additional peaks observed in the bound form are labelled in gold. B) Overlaid 2D ^1^H-^13^C INEPT-HETCOR spectra of aSyn pS129 without DNAJB1 (teal) and with DNAJB1(crimson). Additional peaks observed in the bound form are labelled in crimson.

Next, we characterized the dynamic residues of the pS129 fibrils. Upon phosphorylation of the fibrils, we noticed that the sample became more opaque than unphosphorylated fibrils. The phosphorylation site at S129 was absent in the HSQC. The HSQC spectrum of aSyn pS129 fibrils exhibited lower sensitivity compared to WT fibrils; however, we were able to tentatively assign twenty-three residues based on the assignments from aSyn WT (Figure 4C). The residues near the phosphorylation site, specifically N122, Q134, E126, D119, M127, and A107 showed a significant decrease in intensity (Figure 4C). In contrast, the spectrum of DNAJB1 bound aSyn pS129 fibrils displayed a drastic reduction in the number of peaks, accompanied by an increase in average peak intensity. In the bound pS129 fibrils, we were only able to observe thirteen peaks, indicating stronger binding between DNAJB1 and the pS129 fibrils compared to the WT fibrils. The peaks that were absent in the pS129 aSyn bound sample were residues V118, D119, D121, N122, E123, E126, M127, E130, E131, D135, and E139. This suggests that more C-terminal residues are perturbed in pS129 fibrils than WT fibrils due to DNAJB1 binding. In contrast to the HSQC of the DNAJB1 bound WT fibrils, in which A140 is intense and doubled, the A140 peak in the spectrum of the bound aSyn pS129 fibrils showed a noticeable decrease in intensity along with splitting. Similar to the WT bound sample, we observed new peaks (E_PB1_and V_PB1_) appearing in the HSQC of pS129 fibrils with DNAJB1 which are highlighted around a purple dashed box. Residue type assignments for E_PB1_ and V_PB1_ were obtained using 2D HETCOR and TOCSY experiments (Figure 5B, S9 respectively). Additionally, we observed A_PB1,_ P_PB1_, and G_PB1_ in the HETCOR and TOCSY spectra, which were not detected in the HSQC.

The overlay of the bound and unbound HSQCs of both fibril types indicate the presence of a new conformation in the presence of DNAJB1 (Figure S6 A,B). Unlike the bound WT spectrum, more of the extreme C-terminal residues were absent in the bound pS129 spectrum. In the WT-bound sample, A140 was split but was intense, whereas in the pS129-bound spectrum, it exhibited reduced intensity and splitting. Additionally, glycines in the WT-bound sample showed a significant decrease in intensity, while those in the pS129-bound sample remained relatively unchanged.

From the plot showing the intensity ratio of the peaks in bound to unbound spectra (I_B_/I_UB_) of aSyn WT and aSyn pS129 fibrils, it is evident that the mode of binding to DNAJB1 is different in the phosphorylated fibrils (Figure 4B, D). The primary binding region in both fibril types appears to be within N122-E130 (Figure 6). In aSyn WT fibrils, DNAJB1 affected residues closer to the fibril core, whereas in aSyn pS129 fibrils, interactions with extreme C-terminal residues were more pronounced.

**Figure 6:**
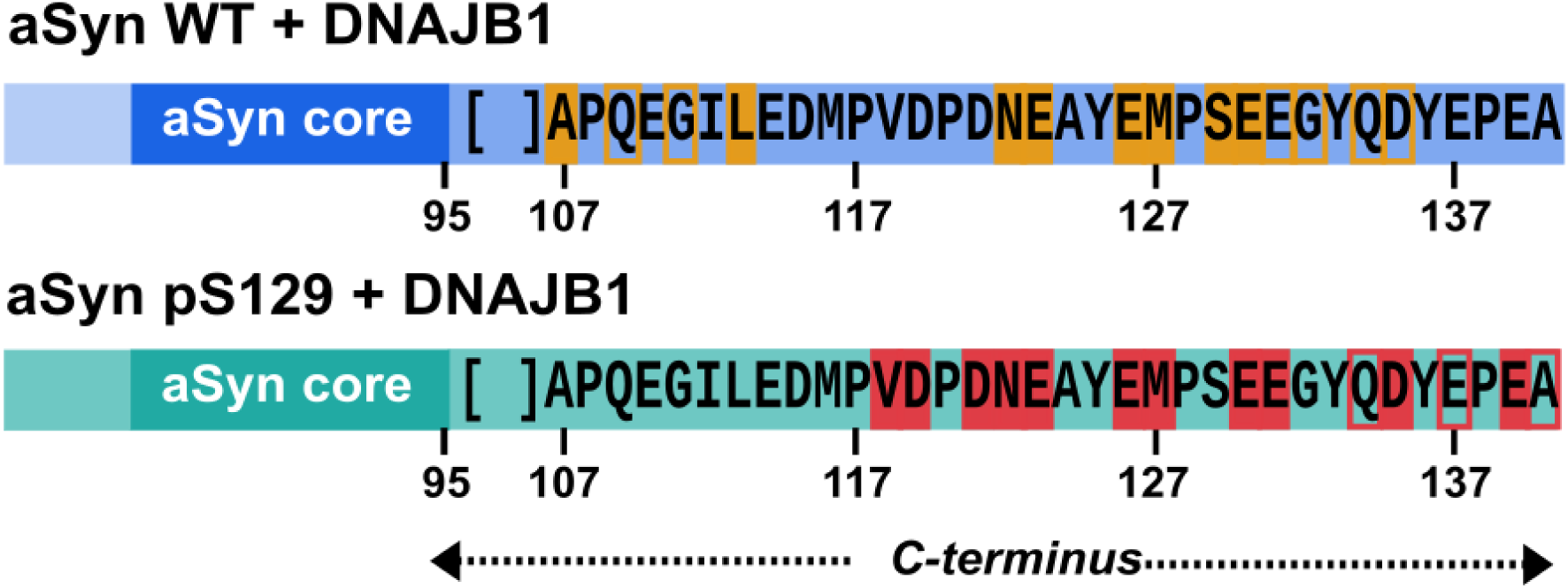
Schematic representation of NMR chemical shift perturbations observed for aSyn WT, and aSyn pS129 fibrils upon DNAJB1 binding. NMR analysis indicates that DNAJB1 interacts with the dynamic C-terminus of aSyn in both fibril types; however, the mode of binding differs. In aSyn WT fibrils, residues affected by DNAJB1 binding are highlighted: those that disappeared are boxed with filled gold rectangles, while those with decreased intensity are boxed with clear gold rectangles. In aSyn pS129 fibrils, residues that disappeared or showed reduced intensity are highlighted with filled crimson rectangles and clear crimson rectangles, respectively. The predominant binding region in both the fibril types appears to be within N122-E130. However, in aSyn WT fibrils, more residues close to the fibril core were affected by DNAJB1, whereas in aSyn pS129 fibrils, more extreme C-terminal residues were affected by DNAJB1.

## Discussion

Our findings highlight the impact of PTMs, particularly in the disordered regions of aSyn fibrils, on chaperone interactions. The aSyn pS129 PTM is especially noteworthy, as 90% of aggregated aSyn in Lewy bodies is phosphorylated at this site.^38^ Our sedimentation assay and solid-state NMR data reveal a preferential binding of DNAJB1 to aSyn pS129 fibrils. This enhanced affinity may be attributed to phosphorylation-mediated electrostatic interactions, which could strengthen chaperone engagement. We also observed a decrease in free monomeric aSyn pS129 compared to the control sample that does not contain DNAJB1, in the SDS-PAGE (Figure 1B). This might be due to pS129 monomer nonspecifically sticking to the fibrils.

Interestingly, while solution-state studies have identified the DNAJB1 binding region on monomeric aSyn as highly specific within the C-terminus (specifically residues 123-129),^27^ our results on aSyn fibrils indicate a more dispersed interaction pattern. In WT fibrils, DNAJB1 interacts broadly along the dynamic C-terminus, affecting both the terminal residues (N122-E130) and also those closer to the fibril core, such as A107 and L113 (Figure 6). This expanded binding site implies that, unlike the specific and localized binding observed in monomeric forms, fibrillar aSyn may present a variety of accessible regions across its disordered C-terminus. The primary binding site is predominantly enriched with acidic residues, consistent with previous reports for monomeric aSyn. In contrast, the secondary binding region extends to V118 which is closer to the fibril core, exhibiting a higher degree of hydrophobicity. While in pS129 fibrils, the DNAJB1 binding region encompasses more of the extreme C-terminus containing more acidic residues (Figure 6).

We observed dramatic changes in our HSQC spectra upon DNAJB1 binding with only a handful of resonances being unaffected. This is surprising for a aSyn:DNAJB1 ratio of 5:1. However, considering that DNAJB1 is a dimer, the binding site to epitope ratio should be rather 2.5:1.

Our analysis shows that several peaks disappear, some get diminished in intensity, and others split into two resonances. In the slow chemical exchange regime and at our aSyn:DNAJB1 ratio, we would expect the spectrum to be dominated by the unbound resonances. In the fast chemical exchange regime, we would not expect to observe any peak splitting. Therefore, we think that the DNAJB1 interaction is in an intermediate exchange regime. In this scenario, the residues at the direct binding interface disappear because the resulting decrease in motion prevents efficient magnetization transfer in the HSQC. However, we do observe peak splitting at the resonances at the edge of the binding site for WT aSyn (e.g. V118, E139, and A140). These residues remain, despite line broadening, dynamic enough to be detected in our J-based spectra but show splitting between the original unbound resonance and the new DNAJB1 bound state.

Notably, as reported by Wintek et al., we also observed variability in chaperone binding depending on fibril preparations (Figure 1C). Differences in binding between WT and pS129 fibrils suggest that PTMs, such as phosphorylation, can modulate chaperone engagement, potentially altering the efficacy of fibril disaggregation. Phosphorylation of aSyn fibrils at S129 may enhance electrostatic interactions with DNAJB1 and expose additional acidic residues in the C-terminus of aSyn pS129 fibrils, thereby expanding the primary binding region. The EM micrographs showed fibril fragmentation in the presence of DNAJB1 (Figure 2), yet there was no significant difference in the fibril width. This suggests that DNAJB1 binding may render fibrils more susceptible to mechanical fragmentation on EM grids, an effect possibly enhanced by phosphorylation at S129. The consistency in fibril width within each fibril type is anticipated, as DNAJB1 binds to aSyn fibrils but lacks the ability to disaggregate them without the assistance of Hsp70 and Apg2^14^.

Future studies should aim to investigate how S129 phosphorylation impacts the disaggregation efficiency of the complete disaggregase machinery, which includes DNAJB1, Hsp70, and Apg2, as this could provide valuable insights. Such studies could reveal whether this post-translational modification enhances or hinders disaggregation and whether the resulting disassembly products differ in toxicity, given that oligomeric fragments are frequently implicated in the pathology of synucleinopathies.

## Supporting Information

Supporting Information: Figures S1-S9

## Author Information

### Corresponding Author

Ansgar B. Siemer - Department of Physiology and Neuroscience, Zilkha Neurogenetic Institute, Keck School of Medicine, University of Southern California, Los Angeles, California, USA;

### Authors

Sayuri Pacheco - Department of Physiology and Neuroscience, Zilkha Neurogenetic Institute, Keck School of Medicine, University of Southern California, Los Angeles, California, USA

Dhanya S. Reselammal - Department of Physiology and Neuroscience, Zilkha Neurogenetic Institute, Keck School of Medicine, University of Southern California, Los Angeles, California, USA

Qingya Zhang - Department of Physiology and Neuroscience, Zilkha Neurogenetic Institute, Keck School of Medicine, University of Southern California, Los Angeles, California, USA

## Author Contributions

S.P. and D.S.R. contributed equally to this work. S.P. and D.S.R. conducted the experiments and wrote the manuscript. Q.Z. performed the SDS-PAGE analyses, and A.B.S. guided the study and revised the manuscript.

## Acknowledgements

This work was supported by the National Institutes of Health under grant numbers R01NS133820 and F31NS131011. We would like to thank Dr. Ralf Langen for providing the plasmid for aSyn. We would also like to thank Drs. Matthew Pratt and Ralf Langen for fruitful discussions.

## For Table of Contents Only

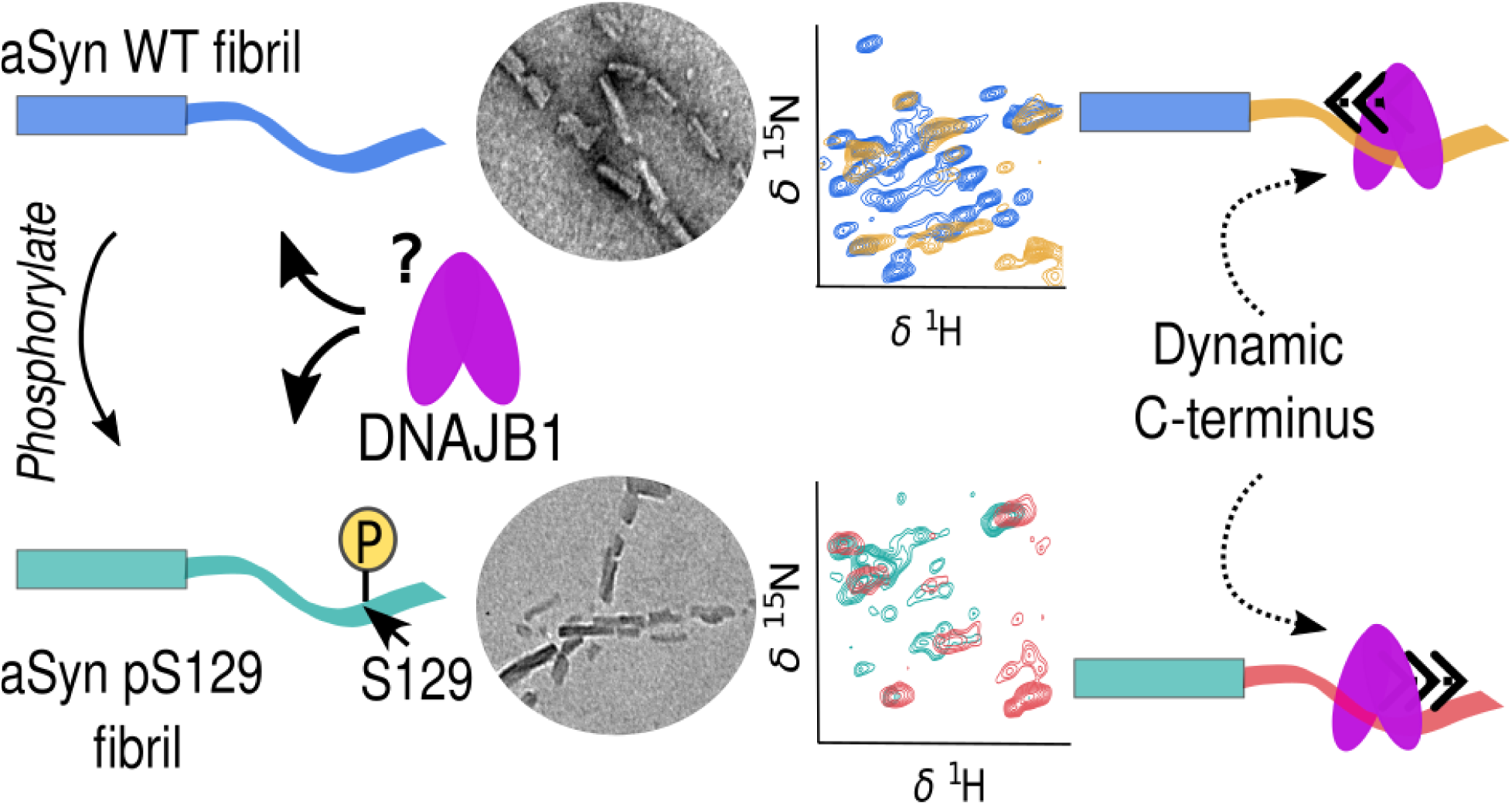

