## Supporting Figures S1-9 for "Phosphorylation of α-synuclein fibrils at S129 changes DNAJB1 binding as probed by solid-state NMR"

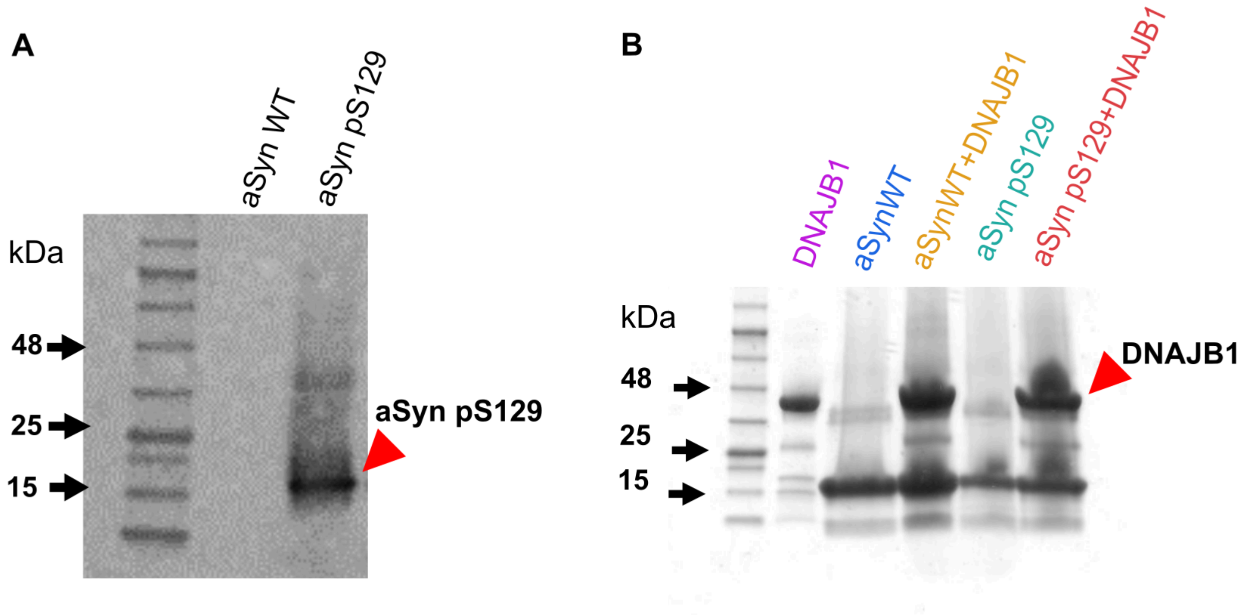


**Figure S1:** A) Phosphorylation of aSyn at S129 by PLK2 (an enzyme specific to this site) was confirmed via Western blot using the Ep1536Y antibody (ab51253), which specifically detects phosphorylated aSyn at S129. Sample from the phosphorylation reaction was solubilized in 0.1% TFA/water and loaded onto lane 3. Thick band near 15 kDa corresponds to aSyn pS129. As a control, solubilised aSyn WT (unphosphorylated) fibrils were loaded onto lane 2. B) SDS-PAGE analysis of the solubilized fibril pellets from the sedimentation assay. Fibril pellets were solubilized in 1% DMSO in TFA/water and loaded onto the gel. Both solubilized fibril samples showed the presence of aSyn and DNAJB1.

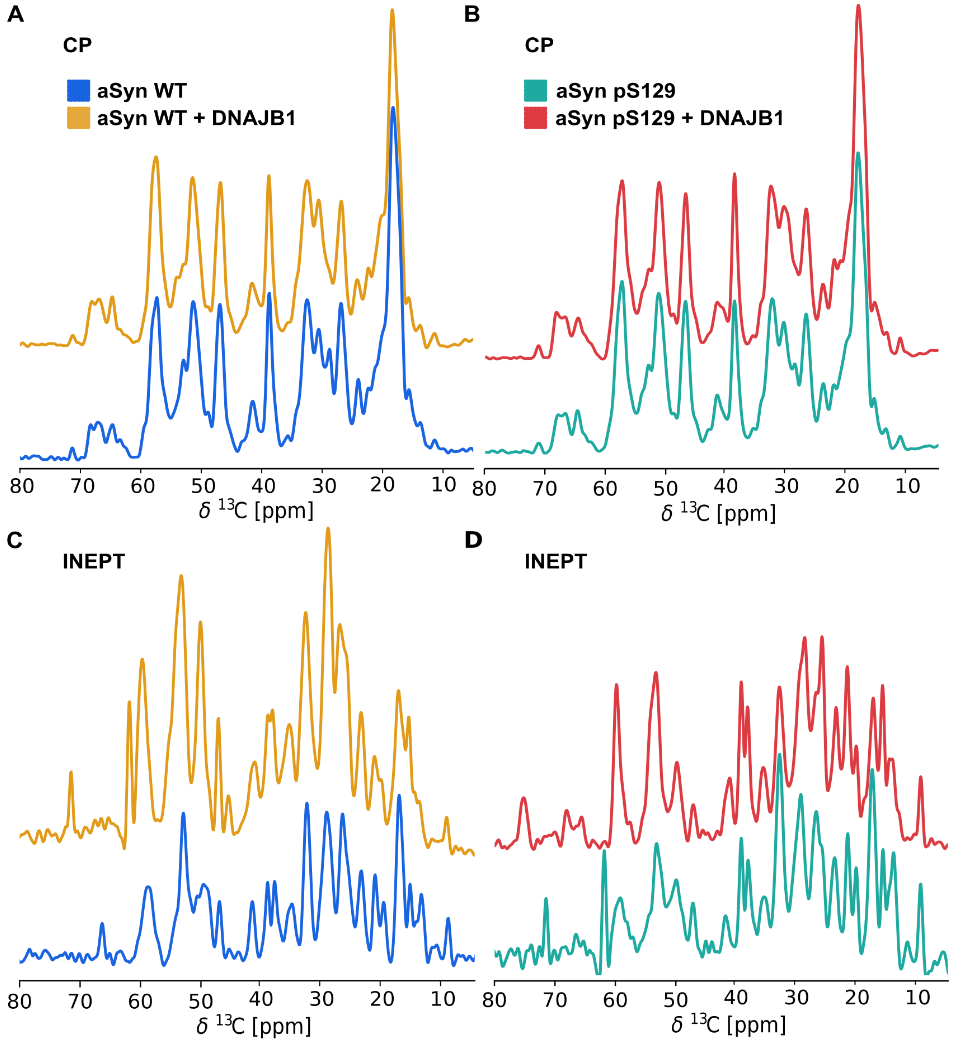


**Figure S2: Overlay of 1D solid-state NMR spectra of aSyn fibrils in the absence and presence of DNAJB1.** A, B) cross polarization (CP) spectra of WT and pS129 fibrils, respectively. Their perfect overlap indicates that the static domains of the fibrils remain unaffected by the addition of DNAJB1. C, D) Insensitive nuclei enhancement by polarization transfer (INEPT) spectra of WT and pS129 fibrils highlight clear changes in the dynamic regions of aSyn fibrils upon binding to DNAJB1. These results suggest that DNAJB1 interaction selectively affects the dynamic regions of aSyn fibrils without altering the static core.

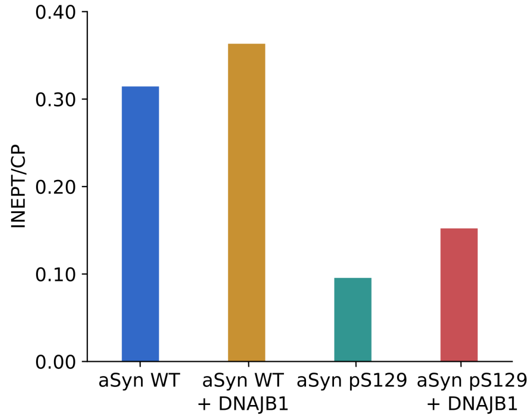


**Figure S3: Dynamics of aSyn WT and pS129 fibrils increase in the presence of DNAJB1.** CP and INEPT spectra of the aSyn WT, and aSyn pS129 were integrated and the INEPT /CP ratio was calculated for each sample. A decrease in the INEPT/CP ratio of the aSyn pS129 fibrils indicates reduced dynamics in the phosphorylated fibrils compared to the WT fibrils. However, in the presence of DNAJB1, both fibril types show an increase in the INEPT/CP ratio, suggesting enhanced molecular dynamics.

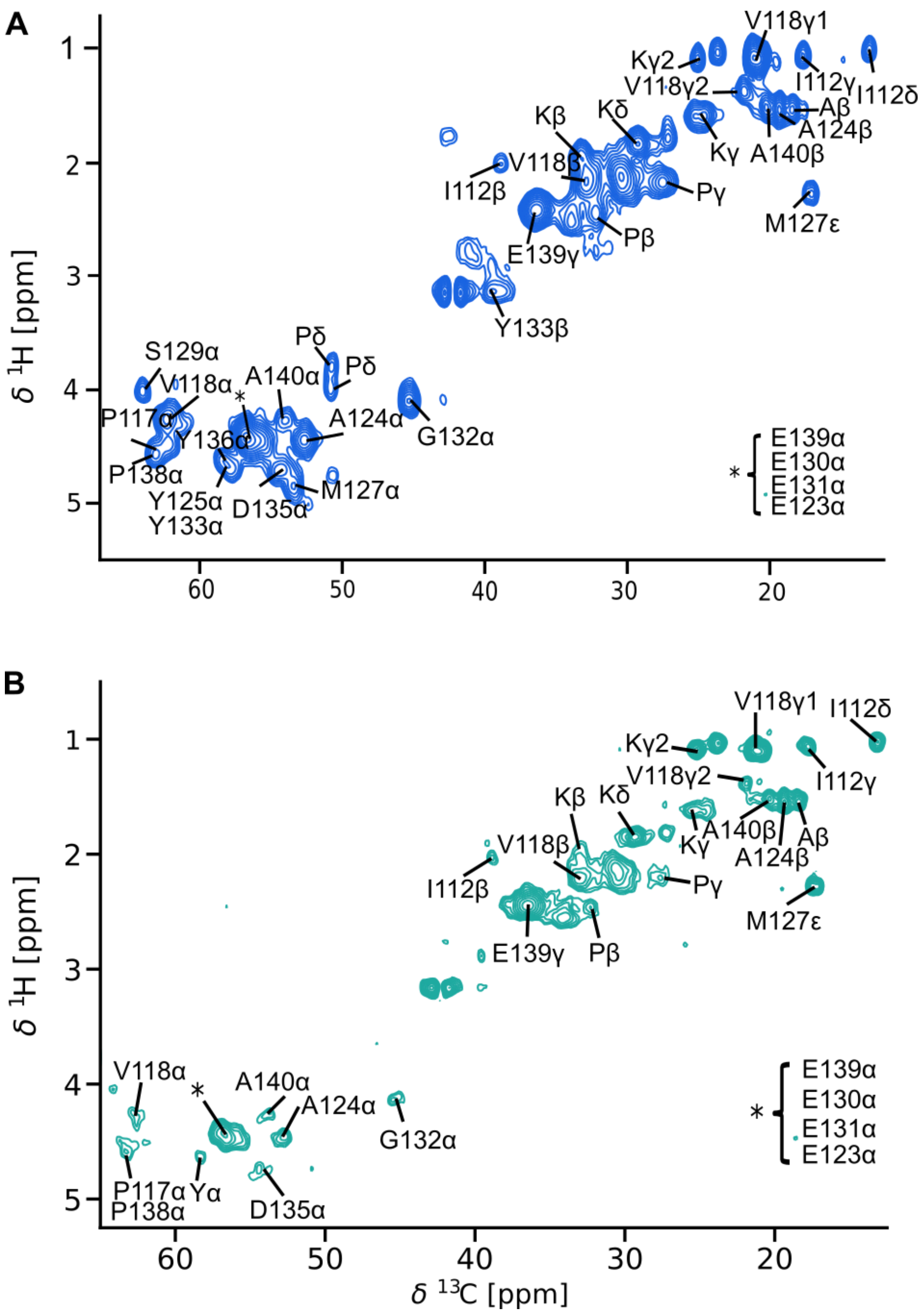


**Figure S4:** $^{1}H$-$^{13}C$ INEPT-HETCOR spectra of aSyn WT fibrils (A), and aSyn pS129 fibrils (B). Site- and residue-specific assignments are indicated.

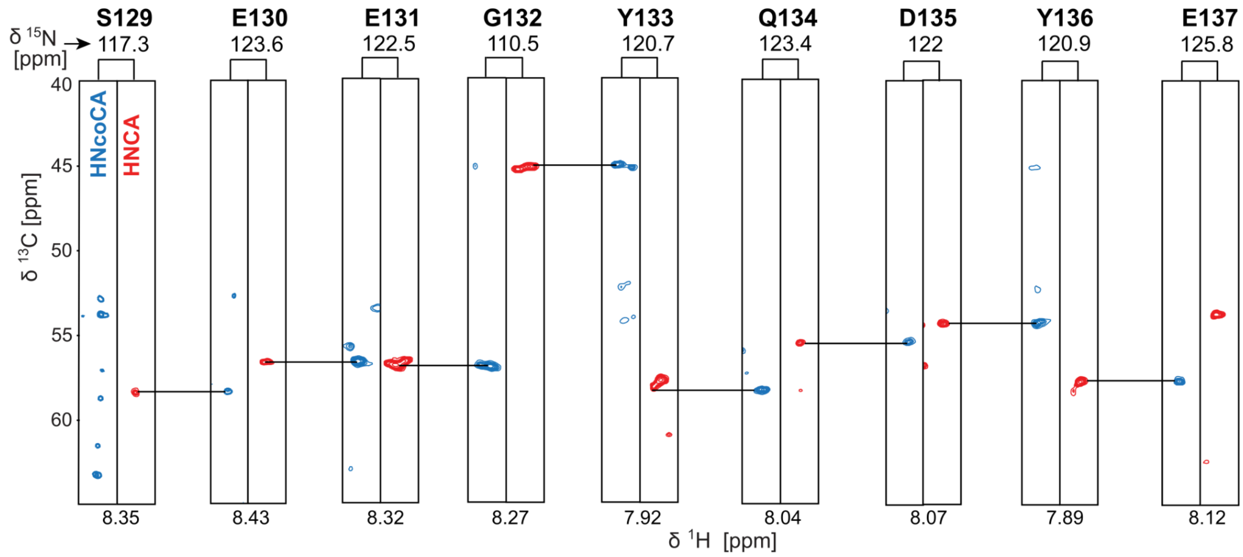


**Figure S5:** Strip plots demonstrating the aSyn WT assignment using HNCA and HNcoCA experiments.

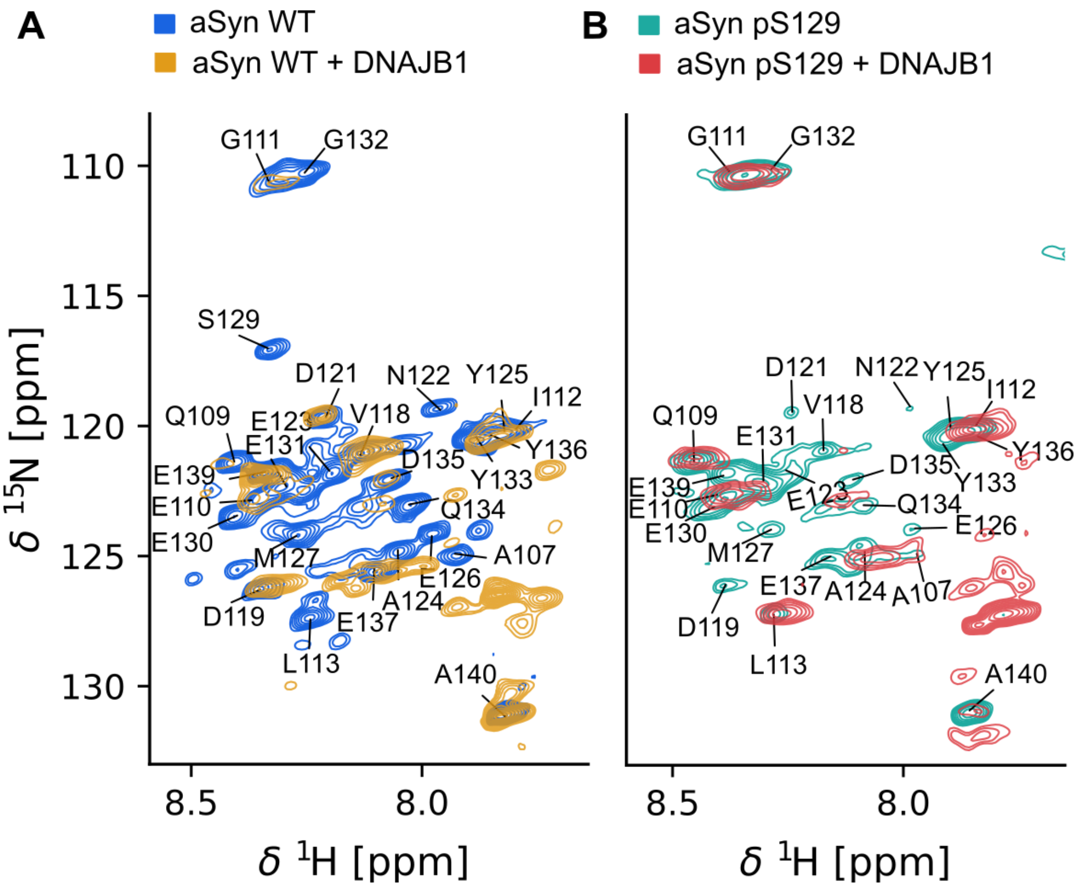


**Figure S6:** Overlaid HSQC spectra of DNAJB1-bound and unbound fibril types: A) aSyn WT, and B) aSyn pS129.

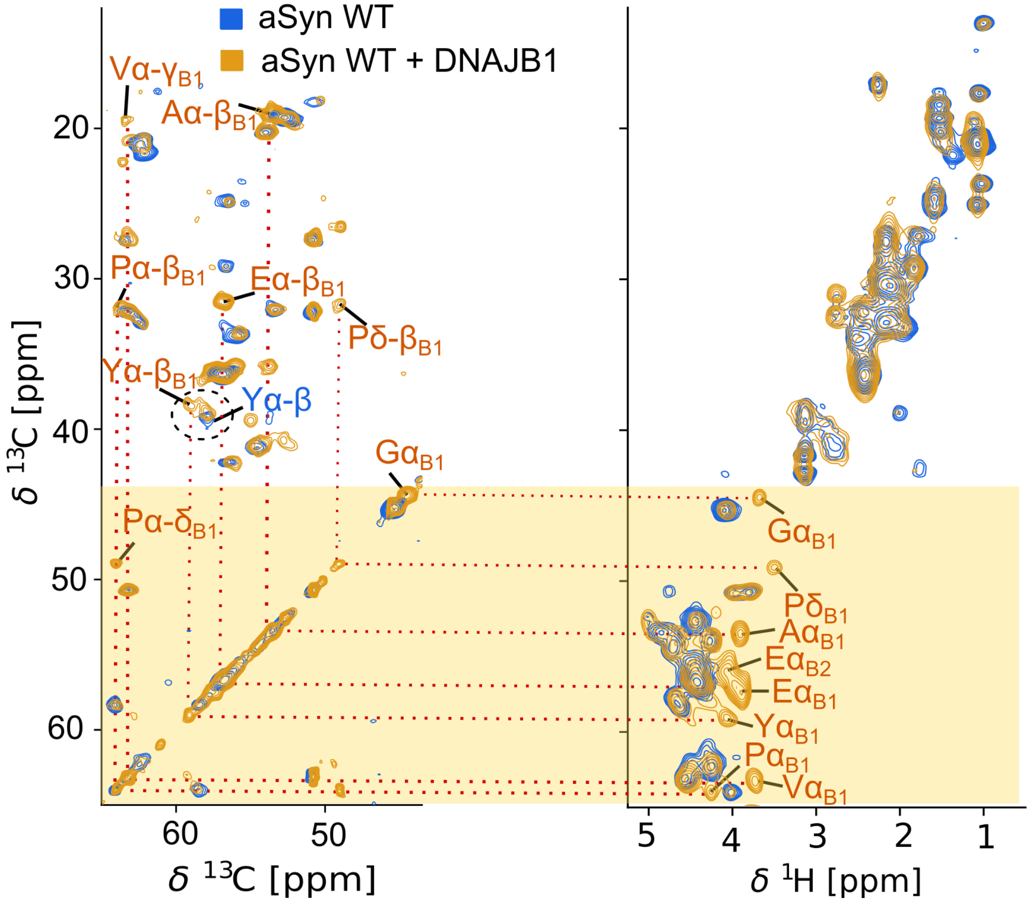


**Figure S7:** Residue-types of the additional peaks in the HSQC and HETCOR spectra of DNAJB1-bound aSyn WT (Fig. 4A and 5A) were assigned using 2D $^{13}C$-$^{13}C$ INEPT-TOCSY (left) and $^{1}H$-$^{13}C$ INEPT-HETCOR (right) experiments. The Cα region of the overlaid spectra, shown in the main Fig. 5A, is highlighted with a yellow-shaded box. Overlaid TOCSY and HETCOR spectra of aSyn with (gold) and without DNAJB1 (blue) display chemical shift perturbations and peak splitting resulting from DNAJB1 binding. Residue types of the additional peaks were identified by correlating the peaks within the spin system.

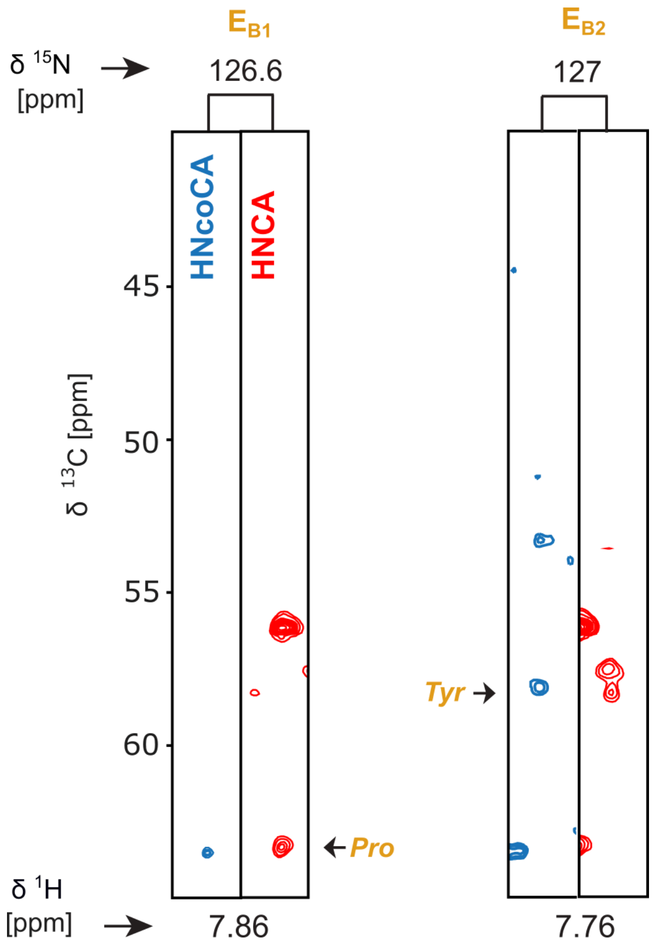


**Figure S8:** The 3D strip planes for the new peaks $E_{B1}$ and $E_{B2}$ from the HNCA and HNcoCA experiments of the DNAJB1 bound aSyn WT fibrils

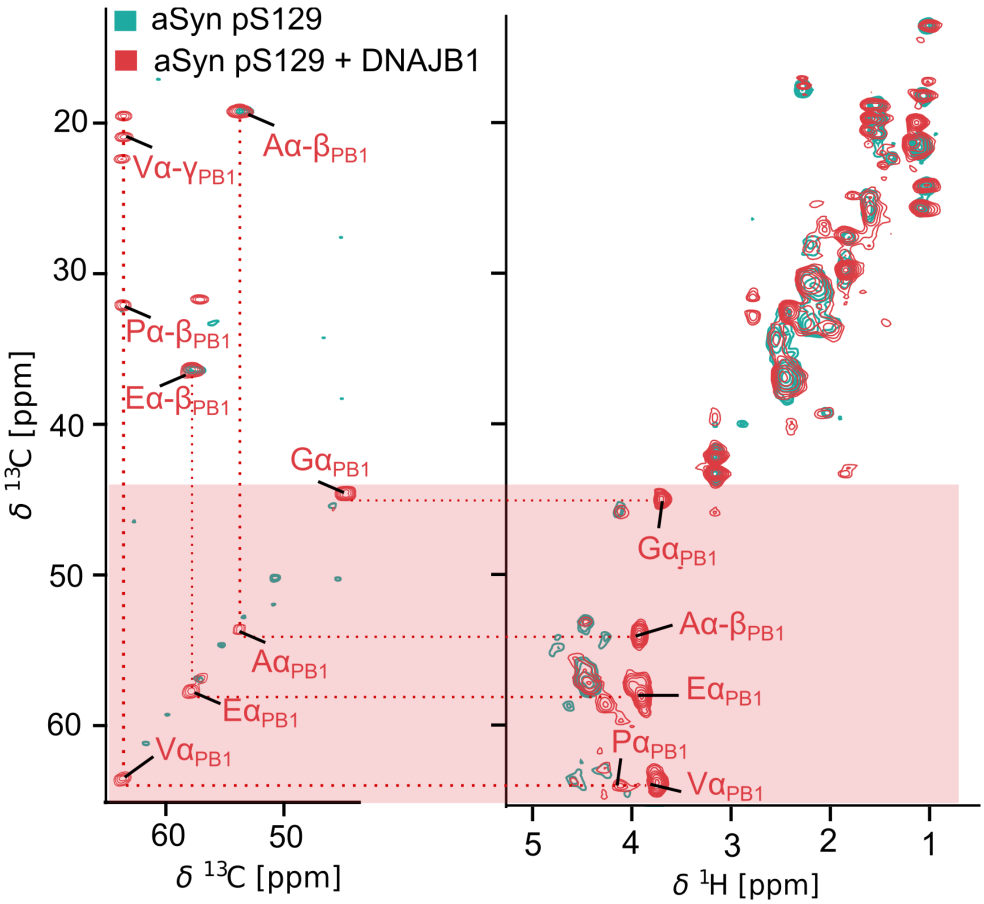


**Figure S9:** Residue-types of the additional peaks in the HSQC and HETCOR spectra of DNAJB1-bound aSyn pS129 (Fig. 4B and 5B) were assigned using 2D $^{13}$C-$^{13}$C INEPT-TOCSY (left) and $^{1}$H-$^{13}$C INEPT-HETCOR (right) experiments. The Cα region of the overlaid spectra, shown in the main Fig. 5 B, is highlighted with a red-shaded box. Overlaid TOCSY and HETCOR spectra of aSyn pS129 with (crimson) and without DNAJB1 (teal) display clear chemical shift perturbations and peak splitting resulting from DNAJB1 binding. Residue types of the additional peaks were identified by correlating the peaks within the spin system.
